# Biodiversity lost: The phylogenetic relationships of a complete mitochondrial DNA genome sequenced from the extinct wolf population of Sicily

**DOI:** 10.1101/563684

**Authors:** Stefano Reale, Ettore Randi, Floriana Bonanno, Valentina Cumbo, Ignazio Sammarco, Antonio Spinnato, Salvatore Seminara

## Abstract

Using next-generation sequencing, we obtained for the first time a complete mitochondrial DNA genome from a museum specimen of the extinct wolf (*Canis lupus*) population of the island of Sicily (Italy). Phylogenetic analyses showed that this genome, which was aligned with a number of historical and extant complete wolf and dog mtDNAs sampled worldwide, was closely related to an Italian wolf mtDNA genome (TN93 and *p*-distances = 0.0012), five to seven times shorter than divergence among Sicilian and any other known wolf mtDNA genomes (distance range = 0.0050 – 0.0070). Sicilian and Italian haplotypes joined a basal clade belonging to the mtDNA haplogroup-2 of ancient western European wolf populations (Pilot et al. 2010). Bayesian calibration of divergence times indicated that this clade coalesced at MRCA = 13.400 years (with 95% HPD = 4000 – 21.230 years). These mtDNA findings suggest that wolves probably colonized Sicily from southern Italy towards the end of the last Pleistocene glacial maximum, when the Strait of Messina was almost totally dry. Additional mtDNA and genomic data will further clarify the origin and population dynamics before the extinction of wolves in Sicily.

## Introduction

During the last few centuries biodiversity has been dramatically destroyed worldwide (WWF 2018). With a few exceptions, range, abundance and genetic diversity of many animal populations declined pressed by the impacts of human population expansion and degradation of natural habitats (Li et al. 2016). In particular, large vertebrates and top predators paid the price of environmental anthropogenic changes. Wolf (*Canis lupus*) is one of the few large predators that managed to survive safely the Pleistocene faunal turnover (Loog et al. 2018). Nevertheless, wolf populations fluctuated widely in the Old and New World through the Pleistocene and more recently, during the last few centuries, in consequence of deforestation, prey overhunting and direct persecution (Leonard et al. 2005; Randi 2011). In the recent past, wolves disappeared from southern and central regions of North America and from most of central European countries. Although wolves are currently recovering, aided by legal protection, controlled hunting and active conservation, the recent demographic declines led to a number of local extinctions, in particular small and isolated populations (Linnell et al. 2008). For instance, the last Honsu wolf (*C. l. hodophilax*), the endemic dwarf subspecies living in three of the main islands of Japan, was killed in 1905 (Ishiguro et al. 2009). The Ezo wolf (*C. l. hattai*), endemic of the Hokkaido, was eradicated from the island by the end of the 1800s (Ishiguro et al. 2010). Recently, also the very small isolated wolf population in the Sierra Morena of central Spain has been definitely eradicated (Gómez-Sánchez et al. 2018).

Fossil remains indicate that wolves were present in Europe at least by the end of the Middle Pleistocene at about 0.5–0.3 million years ago (Sotnikova & Rook 2010). However, those ancestral wolf populations, which showed distinct ecology, morphology and genetics, were completely substituted by contemporary wolves that rapidly spread across all Europe c. 25.000 – 20.000 year ago (Loog et al. 2018). The extant wolves of the Italian peninsula(*C. l. italicus*) are genetically divergent from all the other wolf populations in Europe, likely due to their long-lasting isolation south of the Alps, to historical and recent anthropogenic bottlenecks (Lucchini et al. 2004; Pilot et al. 2014). Wolves in peninsular Italy hardly survived at the end of the 1970’ in the southernmost parts of the Apennine, distributed in two small isolated subpopulations counting less than 100 individuals in total (Boitani 1984). In the last two centuries wolves in the Alps and Apennine harboured much more mtDNA diversity that presently (Dufresnes et al. 2018). Since then, the recovery of the Italian wolves has been spectacular, but at that time wolves living in the island of Sicily already went extinct (Angelici et al. 2018).

Information on the Sicilian wolf population is scanty: its phylogeographic origin, historical distribution and abundance in the island are largely unknown and only partially reconstructed anecdotally. Most probably wolves were already rare in first half of the 1800’, likely due to habitat and natural prey loss, and direct persecution. Crossbreeding with free-ranging dogs was already a threaten (Minà Palumbo 1868). However, purpose hunting has likely been the main cause of the Sicilian wolf population decline and final extinction. Documented reports mentioned the killing of seven wolves in 1891 in San Fratello (Messina) and one wolf in 1902 in San Pietro (Caltagirone) (Pratesi 1978). Although reports mentioned the alleged presence of wolves until 1959, the last documented wolf was killed in 1935 in the woodlands in Ficuzza (Giovanni Giardina and Andrea Milazzo; *pers. com*.). Morphologic and genetic analyses showed that the skin of a canid shoot in 1924 in Bellolampo (Palermo; preserved at the Regional Museum of Terrasini), belongs to a domestic dog or a hybrid (Angelici et al. 2018). A few other wolves killed before 1935 are preserved in museums in Sicily or in peninsular Italy (Angelici et al. 2018).

Wolves in Sicily have been cut of their mainland populations in consequence of Mediterranean sea-level fluctuations and flooding of land-bridge connections. Paleogeographic reconstructions and paleontological data documented periods of intense African-Sicily faunal interchange through the strait of Sicily during the Messinian land-bridge, *c*. 5.3 million years ago (Stock et al. 2008). More recently, concurrently with the Pleistocene glacial maxima, the northeastern coasts of the island were in connection with the southern tips of peninsular Italy (Calabria) (Antonioli et al. 2014). The Messina strait, currently 3.2 km wide and 80-120 m deep, has been repeatedly dried at glacial maxima, and Pleistocene temporary land-bridges have been used by a number of animal populations to colonize the island (Antonioli et al. 2014). Wolf remains have been identified in the fossil record of peninsular Italy since the late Middle Pleistocene, c. 340,000 years ago (Anzidei et al. 2012), but modern wolf populations (*Canis lupus subspp*.) colonized the peninsula much more recently, towards the end of the last glaciations (Bertè and Pandolfi 2014). Although morphological traits and preliminary molecular identifications of a few Sicilian wolves specimens and body remains preserved in museum collections were recently described (Angelici et al. 2018), the origin and phylogenetic relationships of the Sicilian wolves are still largely unknown.

In this study we present a new sequence of a complete mtDNA genome obtained from a stuffed Sicilian wolf specimen, aiming at: 1) evaluating the position of this Sicilian wolf mtDNA genomes within the phylogenetic framework of extant and historical wolves in Italy and worldwide; 2) obtaining reliable estimates of haplotype divergence times using complete mtDNA genomes and not limited to very short mtDNA sequences; 3) identifying the likely origin of the sequenced mtDNA genome, either if from a wolf or a dog ancestral maternal population via hybridization; and 4) contributing to reconstruct a plausible scenario of wolf colonization of the island of Sicily.

## Materials and Methods

### DNA extraction

A single tissue sample (the inner part of a nail) was collected from a stuffed wolf specimen preserved at the Civic Museum “Baldassare Romano”, Termini Imerese (Palermo; Italy). Although some authors assert that the wolf (Fig. 1) was probably killed on Monte San Calogero near Termini Imerese (Palermo) in the last years of the 19th century (Angelici and Rossi 2018), we didn’t found truthful historical data regarding the its origins and the years of acquisition by the museum. The nail surface was decontaminated by UV radiation for 30 minutes. We collected inner dry tissue remains by drilling the nail, and the powder was stored in sterile UV-decontaminated test tubes. DNA was extracted twice using a specific forensic kit and procedure (ChargeSwitch^®^ Forensic DNA Purification Kit, Thermo Fisher). In short: the sample was lysed overnight under agitation at 50°C; the DNA was bind to magnetic beads and then cleaned through three washing step employing a magnetic tube support; finally the DNA was recovered washing the magnetic beads with 50 μl of elution buffer. The DNA quality and concentration was checked and quantified by spectrophotometer analysis using Nanodrop (Thermo Fisher Scientific, Waltham, Massachusetts, USA), Qubit (Thermo Fisher Scientific, Waltham, Massachusetts, USA) and TapeStation (Agilent Technologies Inc. Santa Clara, California) equipments. To minimize risk of contaminations by exogenous DNAs, all sample manipulations were done in a DNA-free area that was never used before for DNA extractions. All bench-tops and equipments were flamed or cleaned with bleach, and ethanol and UV irradiated for 60 minutes before and after their use. We used pipette tips with aerosol filters. The DNA library was prepared in a DNA-free area using decontaminated equipments only.

**Figure 1.**
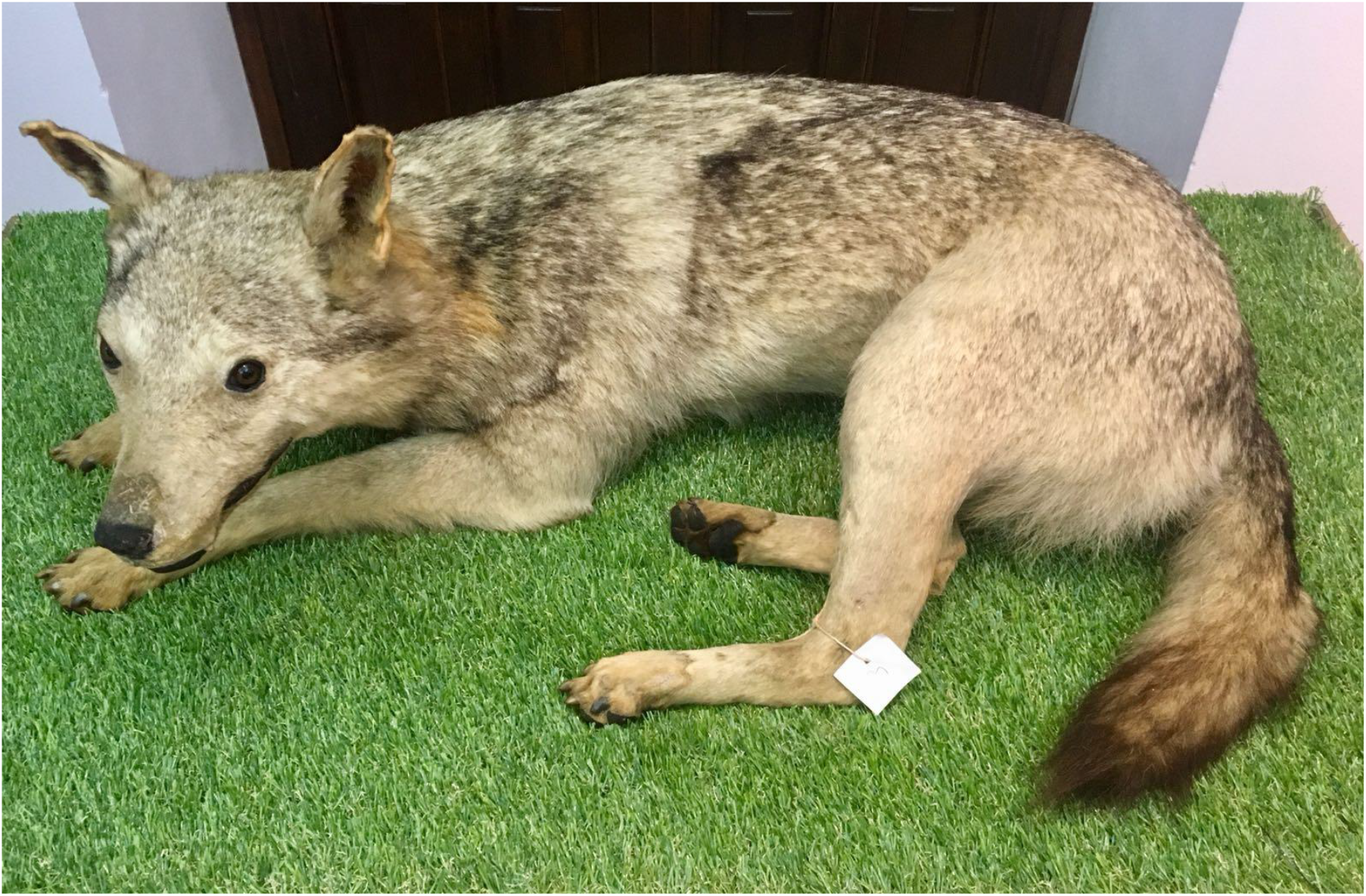
A picture of the sampled and genotyped Sicilian wolf from the Civic Museum “Baldassare Romano”, Termini Imerese (Palermo; Italy).

### NGS library preparation and DNA sequencing

We used the Illumina Truseq DNA Nano kit (Illumina Inc., San Diego, CA, USA) for library preparation according to the manufacturer instructions with some modifications. We did not perform the fragmentation step due to the small size of the input DNA as shown by TapeStation spectrophotometer analysis. We performed end repair to blunt the DNA fragments on a thermal cycler at 30°C for 30 minutes using the End-Repair Mix2. Appropriate library size for sequencing was selected removing the shortest DNA fragments with 250 μl of undiluted Sample Purification Beads reagent (SPB). A single adenine nucleotide was added to the 3’ ends of the blunt fragments to prevent them from ligating to each other and to provide complementary bases for adapters. In detail: 12.5 μl of A-Tailing Mix were added and the following thermal cycler program was used: 37°C for 30 minutes, 70°C for 5 minutes, 4°C for 5 minutes, hold at 4°C. Adapter ligation was performed adding 2.5 μl Resuspension buffer, 2.5 μl LIGation Mix2, 2.5 μl DNA adapter solution, and incubating at 30°C for 10 minutes. The reaction was stopped adding 5 μl of Stop Ligation Buffer. Adapter dimers were removed from the library cleaning the ligated fragments by SPB. The library was PCR-amplified with 5 μl of PCR Primer Cocktail that anneals to the ends of the adapters and 20 μl of Enhanced PCR Mix at 95°C for 3 minutes, 8 cycles of: 98°C for 20 seconds, 60°C for 15 seconds, 72°C for 30 seconds, 72°C for 5 minutes, hold at 4°C. The amplified DNA was cleaned with SPB. Accurate quantification of DNA libraries, a critical step to produce optimal cluster densities across every lane of the flow cell and achieve the highest quality sequencing data, was assayed using a fluorometric method based on dsDNA binding dyes. The distribution of the amplified DNA fragments size was centered at the 295 bp. To obtain the input DNA solution for the sequencing reaction on Illumina Cartridge, the selected size of DNA was at first normalized at 4nM and then diluted at 12.5 pM. The sequencing step was performed with a Illumina MiSeq sequencer using a SBS MiSeq Reagent Kit v2 (Illumina). PhiX Control library (v2; Illumina) was added to the library. The libraries were sequenced with a 150 Paired End MiSeq run. Image analysis, base calling and data quality assessment were performed on the MiSeq and the BaseSpace cloud software was used to generate the final FASTQ files.

### Bioinformatic analyses and mtDNA genome assemblage

The output files from the Illumina sequencer were pre-processed removing adapter sequences, low quality and short read sequences using AdapterRemoval (Schubert et al. 2016) and Cutadapt (Martin 2011). Edited paired-end reads were merged with their overlapping regions using PEAR (Zhang et al. 2014). The processed reads were aligned using an Italian wolf complete mtDNA reference genome (GenBank accession number KF661048) and the BWA-MEM aligner algorithm (Li and Durbin 2010). Mapped reads in BAM format were filtered using MapDamage 2.0 (Jónsson et al. 2013) and quality scores of C->T or G->A transitions, potentially due to post-mortem DNA damage, were recalculated according to the position in reads and damage patterns. Filtered reads were extracted from BAM file using SAMTools 1.4 (Li et al. 2009) and then assembled into a complete consensus sequence with Spades 3.11 (Bankevich et al. 2012). The quality of assembled mtDNA genome was evaluated using Quast tool (Gurevich et al. 2013). Male-specific DNA sequences from the sex-determining region Y protein (SRY gene) were extracted from the reads of the Sicilian wolf genomes by means of Bowtie 2.3.4.1 (Langmead et al. 2012), using a dog sequence (AF107021.1) as reference. To confirm our sequences, we blasted (Altschul et al. 1990) the extracted reads, which had an average length of 120-130 bp, against GenBank database.

### Phylogenetic analyses and estimates of divergence times

We used ClustalW (Higgins et al. 1994) in MEGA X (Kumar et al. 2018) to align the new Sicilian wolf mtDNA concatenated genome with the following mtDNA genomes downloaded from GenBank:

– Set#1: the complete wolf mtDNA genomes (including the control-region; CR) used by Koblmuller et al. (2016), including three historical wolf samples, 14 dog (*C. l. familiaris*) genomes and five Himalayan wolves (named *C. l. laniger* or *C. l. chanco*) used as outgroups;
– Set#2: the modern wolf mtDNA genomes (CR excluded) used by Thalmann et al. (2013), including four coyotes (*C. latrans*) used as outgroups;
– Set#3: a subset of wolf mtDNA genomes (CR excluded) used by Panget al. (2009) and Matsumura et al. (2014), including two historical Ezo wolf (*C. l. hattai*) and five historical Honsu wolf (*C. l. hodophilax*); four coyote mtDNA genomes were used as outgroups.

The alignments were manually checked and adjusted. The control-regions, which were incompletely sequenced in some genomes, were extracted and aligned separately with partial sequences by Dufresnes et al. (2018) obtained from museum specimens originating from the historical wolf populations which lived in the Alps, Italian peninsula and Sicily. Moreover, the mtDNA CR were blasted in GenBank to search for eventual matching with dogs or other wolf CRs. The Sicilian wolf mtDNA protein coding genes were extracted, concatenated and aligned with the homologous coding gene sequences of the three Italian wolf mtDNA genomes in GenBank: KF661048.1 (Thalmann et al. 2013); KU696389.1 and KU644662.1 (Koblmuller et al. 2016). Short overlapping DNA segments, eventual incomplete stop codons and mutations at the three codon positions were identified using MEGAX. We analysed the Set#1 and Set2# alignments using neighbor-joining procedure (NJ; Saitou and Nei 1987) implemented in MEGAX, with the TN93 model (Tamura and Nei 1993), assuming heterogeneous gamma distribution of mutation rates (*a* = 0.5) and heterogeneous lineages evolution. All positions containing gaps, including a variable copy number repeat unit in the CR (about 511 bp from nucleotide 16,040 to 16,550 in the reference dog sequence NC002008), and missing data were pairwise-deleted in the analyses. Support to the phylogenetic tree internodes were determined by 1000 interior-branch length test of minimum evolution trees (ME; Nei and Kumar 2000) and by 10,000 bootstrap samplings of NJ trees (Tamura et al. 2013). Four coyote mtDNA sequences: DQ480509, DQ480510 and DQ480511 (Björnerfeldt et al. 2006); KF661096 1 (Thalmann et al. 2013) were used as outgroups. Bayesian phylogenetic trees were obtained using BEAST 2.5.1 (Drummond et al. 2012), with the HKY+G model (Hasegawa et al. 1985). Markov chain Monte Carlo (MCMC) samples were drawn every 1000 generations from a total of 1,000,000 generations, following a discarded burn-in of 100,000 generations. BEAST was used for age estimates of the nodes of phylogenetic trees. The HKY+G model was assumed for the nucleotide substitution. Both the uncorrelated log-linear model (Drummond et al. 2006) and the strict clock model were tested for the molecular clock, and both the Bayesian skygrid model (Gill et al. 2013) and a constant model for the population size. The convergence and performance of different models were assessed using Tracer 1.7.1 (Drummond and Rambaut 2007). A 10% of the MCMC generations were discarded as a burn-in. BEAST was used to infer the age of the most recent common ancestor (MRCA) of a clade joining the Italian and Sicilian wolf mtDNA, which was calibrated using the divergence times among Japanese wolves (MRCA = 46,800; 95% highest probability density HPD = 37,500–58,000 years) as estimated by Matsumura et al. (2014). The trees were visualized with FigTree 1.4.4, or with TreeAnnotator in BEAST.

## Results

We successfully extracted good-quality DNA from the inner nail tissue of an ancient Sicilian wolf museum specimen. The quantity and quality of the extracted DNA was good enough to obtain reliable sequences, which allowed reconstructing a complete mtDNA genome by next-generation sequencing procedures on Illumina platform. DNA quality was preliminary assessed to ensure successful NGS results. A Nanodrop value of the ratio 260/280 nm = 1.9 indicated a low presence of inhibitors; a Q-bit spectrophotometer analysis showed a total double-strand DNA concentration of 1.6 ng/μl. A TapeStation capillary electrophoresis led to visualize a normal distribution of the DNA fragments, ranging from 40 to 1040 bp and centered at 250 bp, with a concentration of 0.6 ng/μl (Fig. 2a). These quality-controls suggested to skip an initial fragmentation step of input DNA for library preparation, and the DNA was directly PCR-enriched with fragments spanning from 165 to 655 bp, and centered at 295 bp, as showed by TapeStation results (Fig. 2b). The reads obtained from the input DNA, which concentration was 2.49 ng/μl, were assembled into a complete Sicilian wolf mtDNA. The length of this mtDNA genome (GeneBank accession number: MH891616.1) was 16 678 bp.

**Figure 2.**
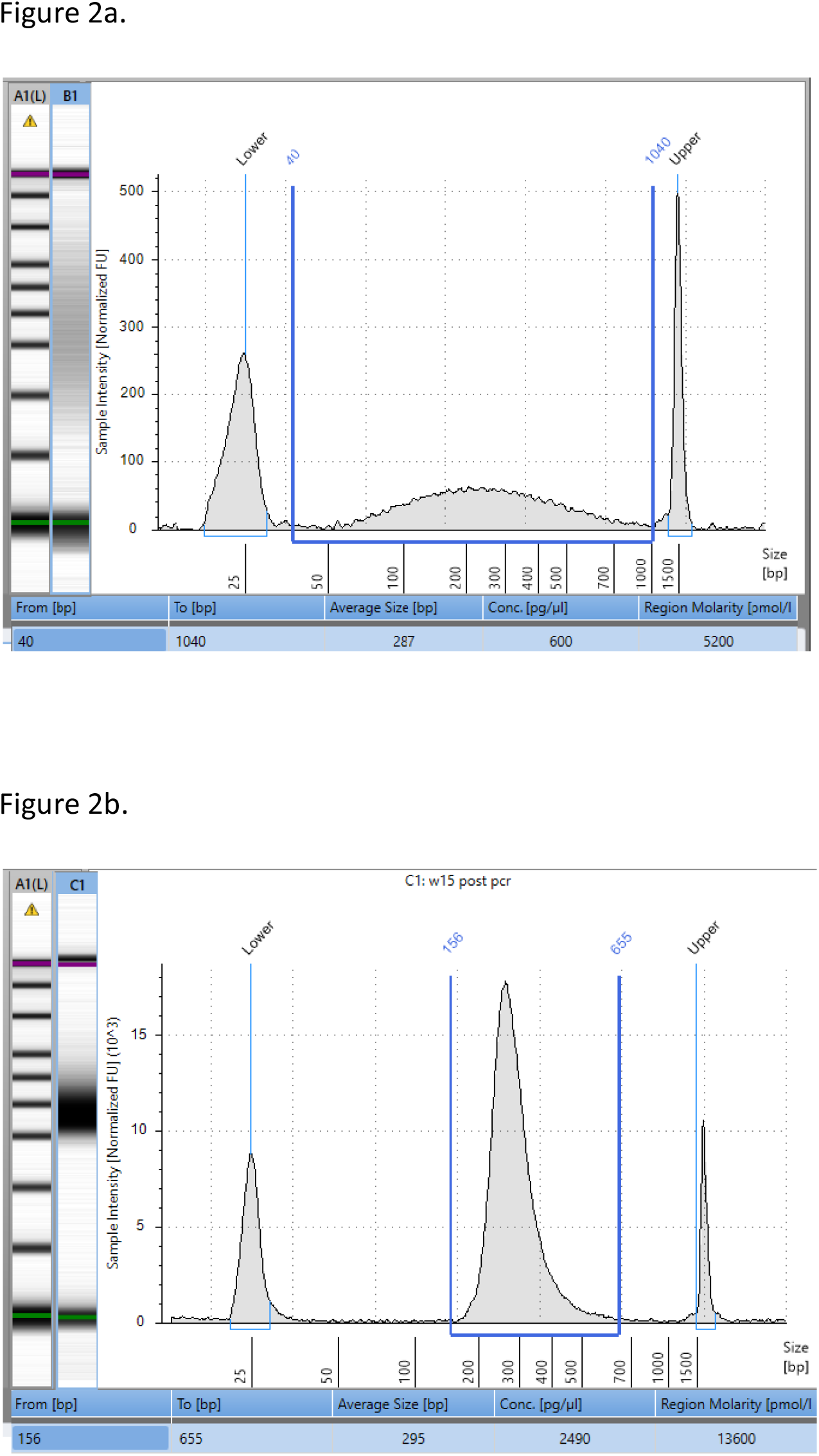
(a) TapeStation spectrophotometer analysis of the extracted DNA from the Sicilian wolf sample. On the left the ladder and the sample run on gel visualization. On the right the fragment size composition of the extracted DNA, spanning from 40 to 1040 bp, with a 287 bp average size at 600 pg/μl concentration. (b) TapeStation analysis of the Sicilian wolf DNA after PCR-amplification. On the left the ladder and the sample run on gel visualization. On the right the fragment size composition of the amplified DNA spanning from 156 to 655 bp, with a 295 bp average size at 2490 pg/μl concentration.

This sequence aligned with reference dog and Italian wolf homologous sequences. All the tRNA, rRNA and protein-coding genes were correctly identified and mapped; these sequences did not show any anomalous stop codon and translated into the expected RNAs or proteins. Thus, we assumed that this mtDNA genome was authentic. In comparison with the known Italian wolf mtDNA genomes, the Sicilian wolf exhibited 14 silent transition substitutions and only one first-position G-A mutation that changed a V into an M aminoacid residue at codon 21 of the ATP6 gene in the Sicilian wolf. The mtDNA CR of the Sicilian wolf was 1219 bp, that is 50 bp shorter than the corresponding sequence of the Italian wolf CR due to 10 missing copies of a CGGTACACGT repeat (Kim et al. 1998). The *p*-distance between the complete mtDNA genome of Sicilian and Italian wolves was *D* = 0.0012 (almost identical to the TN93 distance), that is five to seven times shorter than *Ds* among the Sicilian and any other wolf mtDNA genomes (*D* = 0.0050 – 0.0070). The *p*-distance between Sicilian and Italian wolves was *D* = 0.0181; hence, as expected, the CR evolved about 10 times faster than the RNA and protein-coding mtDNA sequences.

We analysed the Set#1 (Fig. 3) and Set#2 (Fig. 4) mtDNA genomes by the NJ and ME procedures in MEGAX. The mtDNA genome of the Sicilian wolf always joined a basal phylogenetic clade that included the Italian wolf and two mtDNA genomes respectively sequenced from a wolf sampled in Poland (KF661045.1) and in Belarus (KU696390.1). This clade (hereafter named the Italian clade) was basal to all the other modern wolf and dog genomes, including the Japanese wolf clade (*C. l. hodophilax*; Matsumura et al. 2014). The Sicilian wolf genome was basal to the peninsular Italian wolf mtDNAs. Bootstrap and interior-branch length tests showed that the Italian clade was 99% – 100% supported.

**Figure 3.**
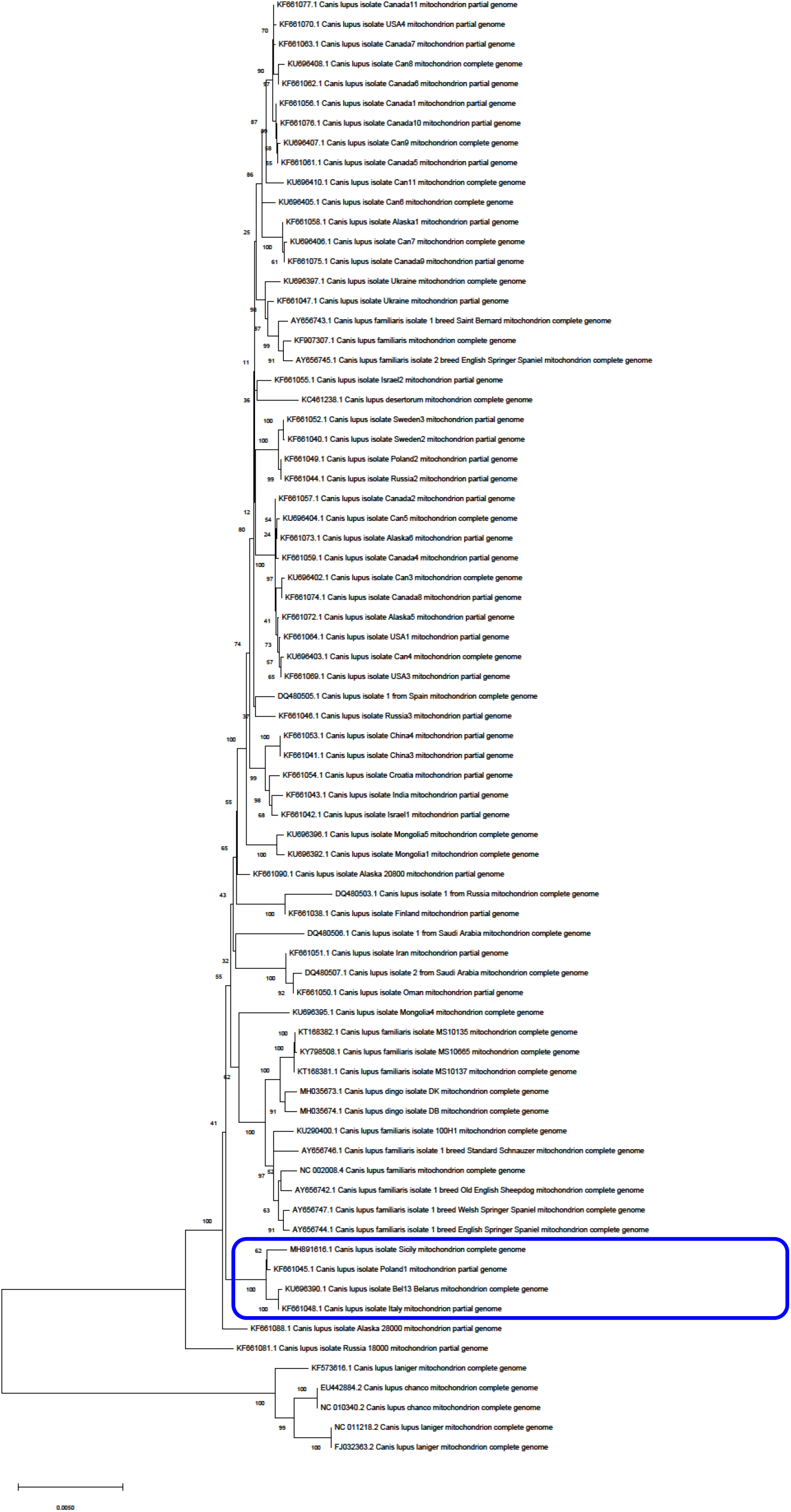
Neighbor-joining tree of complete wolf mtDNA genomes used by Koblmuller et al. (2016), including three historical wolf samples, 14 dog (*C. l. familiaris*) genomes and the new Sicilian wolf mtDNA genome. Five Himalayan wolf (here named *C. l. laniger* or *C. l. chanco*) mtDNAs were used as outgroups. The Italian clade is indicated. Bootstrap values at the internodes.

**Figure 4.**
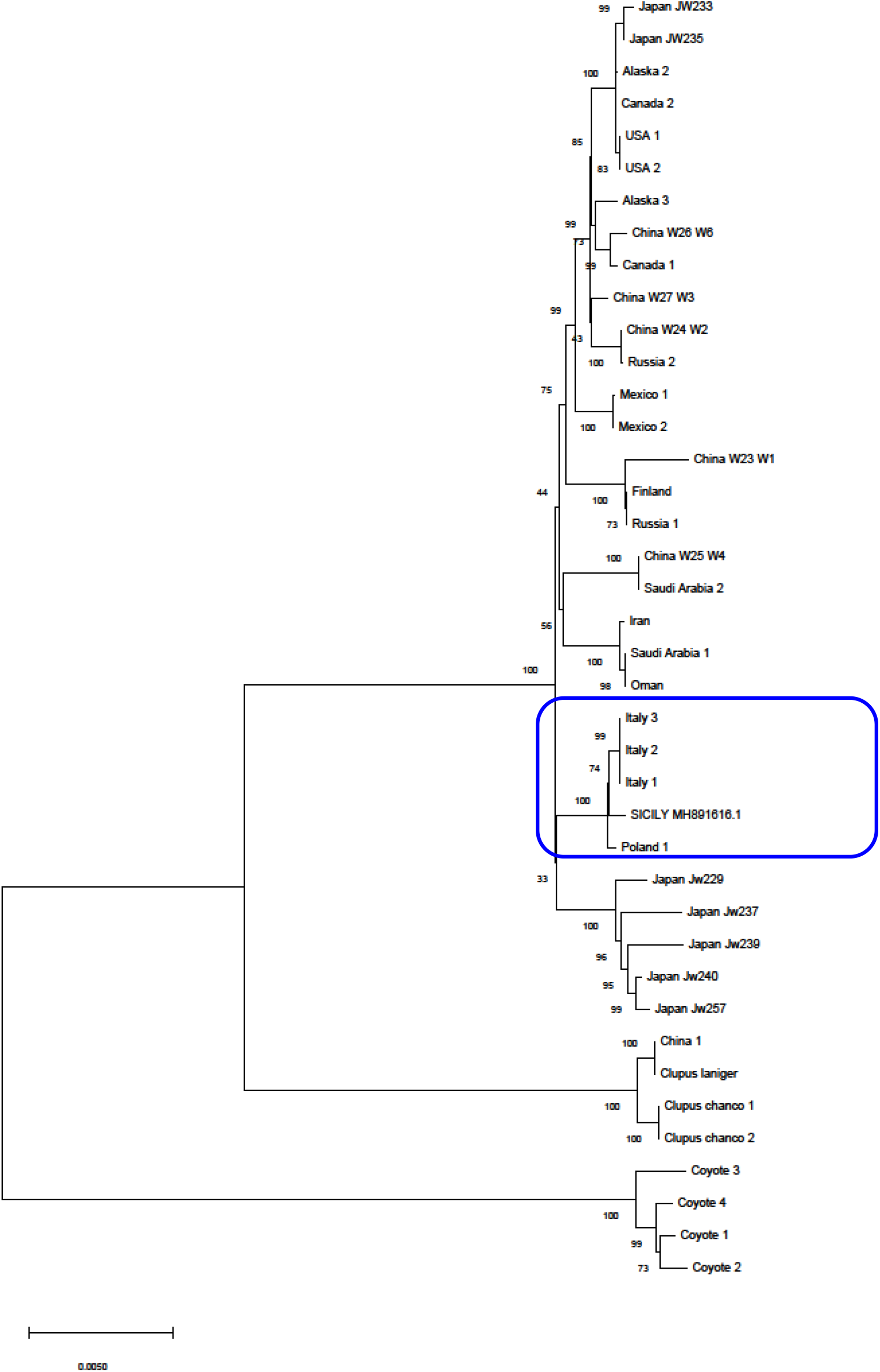
Neighbor-joining tree of modern wolf mtDNA genomes (control-region excluded) used by Thalmann et al. (2013), and the new Sicilian wolf mtDNA genome. Four coyote (*C. latrans*) mtDNAs were used as outgroups. The Italian clade is indicated. Bootstrap values at the internodes.

The Bayesian consensus phylogenetic trees (Fig. 5) obtained analysing the Set#3 mtDNA genomes fully supported the MEGAX results. The Japanese and Italian wolf clades were basal to all the other modern wolf and dog genomes. The average coalescence time of the Italian clade, as estimated in BEAST following Matsumura et al. (2014), was MRCA = 13,400 years (with 95% highest probability density HPD = 4000 – 21,230; Fig. 6). The divergence times of the Italian wolf and Japanese wolf from the other wolf clades were similar (*c.* 100,000 years), further highlighting the ancient origins of wolves in peninsular Italy and Sicily in comparison to the other wolves and dogs worldwide. The Italian and Sicilian wolf mtDNA haplotypes belong to the wolf haplogroup-2, that includes all the ancient wolves sampled in western Europe dating from between 44,000 and 1200 years ago (Pilot et al. 2010).

**Figure 5.**
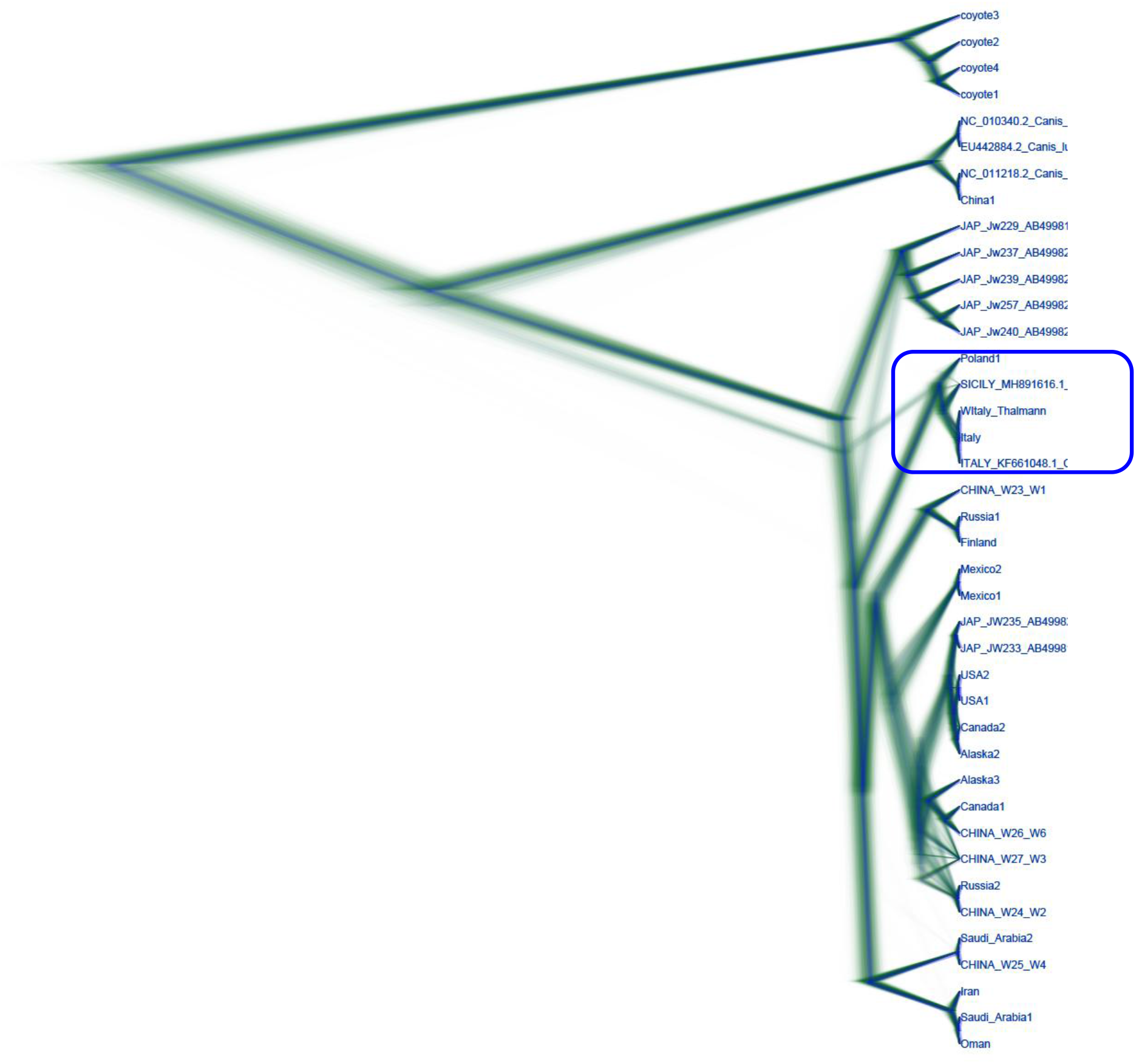
Consensus Bayesian phylogenetic tree computed by BEAST 2.5.1 (Drummond et al. 2012) with the HKY+G model (Hasegawa et al. 1985). We used a subset of wolf mtDNA genomes (control-region excluded) published by Pang et al. (2009) and Matsumura et al. (2014), including two historical Ezo wolf (*C. l. hattai*), five historical Honsu wolf (*C. l. hodophilax*) and the new Sicilian wolf mtDNA genome. Four coyote mtDNAs were used as outgroups. The Markov chain Monte Carlo samples were drawn every 1000 generations from a total of 1.000.000 generations, following a discarded burn-in of 100.000 generations.

**Figure 6.**
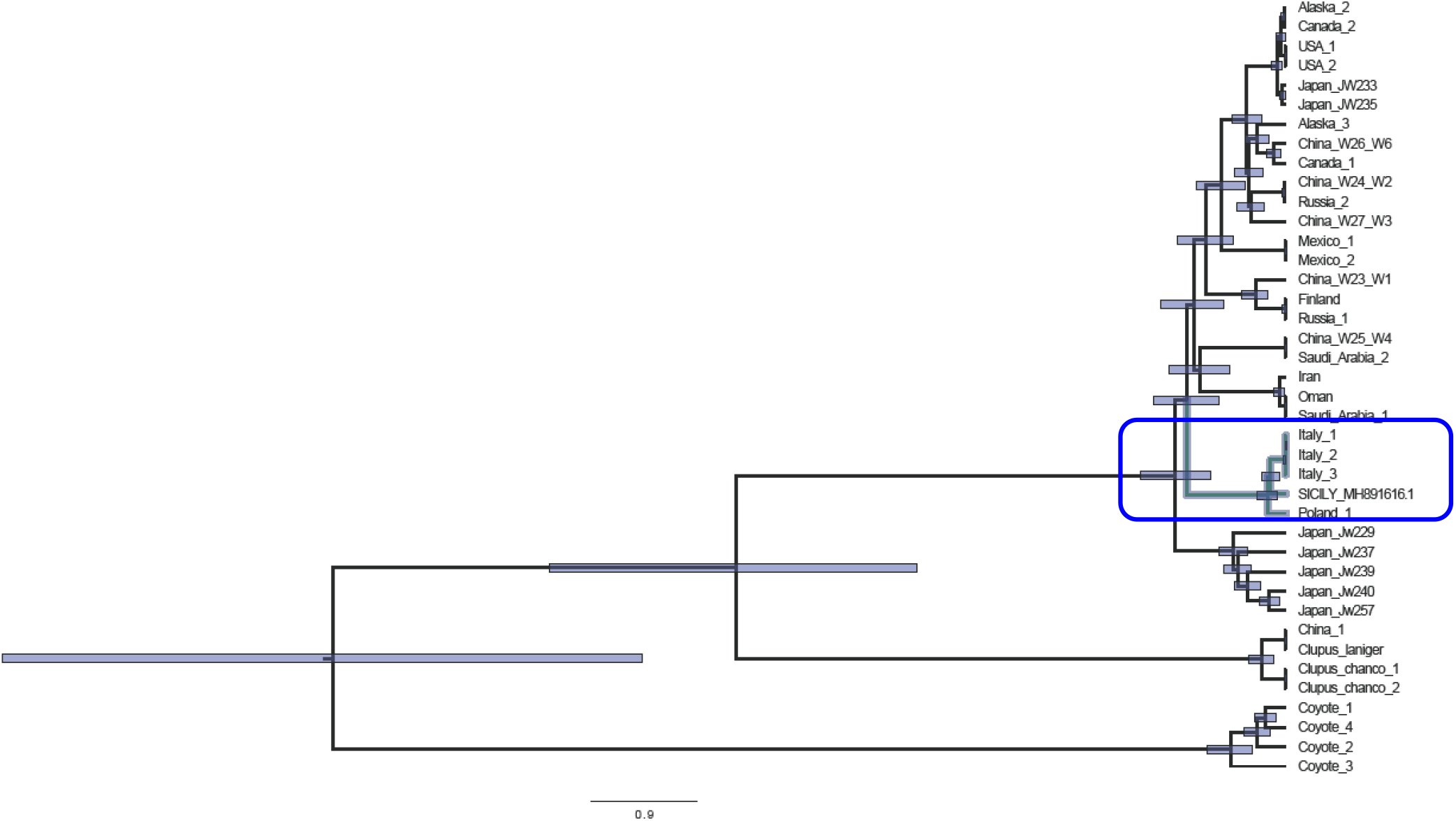
Age estimates (indicated by bar lengths) of the nodes of the consensus phylogenetic trees computed by BEAST 2.5.1 with the HKY+G nucleotide substitution model.

The mtDNA CR of the Sicilian wolf was blasted in GenBank to search for eventual matching with domestic dog CRs. The CR of the Sicilian wolf did not match with any of the dog sequences known so far, thus supporting its origin in a wild wolf populations. The sequence was identical to a partial mtDNA CR sequence found by Dufresnes et al. (2018) in a different wolf sample from Sicily (their haplotype H3 from sample AN855; Museo di Zoologia P. Doderlein, Palermo). Thus, this mtDNA genomes was apparently unique for the extinct wolf population of Sicily.

We analyzed the stored genomic DNA reads to search for specific sex markers. We identified SRY sequences matching with the homologous *Canis lupus* chromosome Y genomic sequences present in GenBank (AF107021.1), thus indicating unequivocally that the studied specimen was a male.

## Discussion

In this study we obtained for the first time a complete mtDNA genome of a Sicilian wolf by NGS technologies. This wolf was likely killed on Mt. San Calogero, near the city of Termini Imerese, in the last years of the nineteenth century, very near to the extinction of the island population, for the last documented wolf was killed in 1935. The control-region of this mtDNA is identical to a partial CR sequenced from a different Sicilian wolf sample (Dufresnes et al. 2018), and is apparently very closely related to a partial CR sequence obtained from another Sicilian wolf specimen, as mentioned by Angelici et al. (2018; although at the moment this sequence is not stored in GenBank). These results suggest that, during the last few decades before the extinction, the wolf population of Sicily showed (at least) two distinct but very closely related mtDNA haplotypes. Phylogenetic analyses also indicate that the mtDNA genome of the Sicilian wolf is closely related to the predominant mtDNA genome of the past and extant wolf population in peninsular Italy (Dufresnes et al. 2018; Randi et al. 2014). The Sicilian and Italian wolf mtDNAs join in a strongly supported clade (the Italian clade) which includes also two mtDNAs sequenced from two wolves sampled in Poland and in Belarus, respectively. The Italian clade is basal to all the other modern wolf and dog haplogroups sequenced so far, with the exception of most of the ancestral sequences obtained from historical wolves (Thalmann et al. 2016; Koblmuller et al. 2016), and from the now extinct Japanese wolves (*C. l. hodophilax*; Matsumura et al. 2014). The origin and fate of the Japanese wolves has been described Matsumura et al. (2014). Both the Japanese and Italian wolf clades, which apparently split *c.* 130,000 – 100,000 years from all the other modern wolf haplogroups worldwide, belong to the mtDNA haplogroup-2 (Pilot et al. 2010). This haplogroup has been detected in the ancient western European wolf population that were largely substituted by the recent spread of modern wolves, which showed the more recent mtDNA haplogroup-1. However, the mtDNA genomes clearly indicate that both extant wolves in Italy and extinct wolves in Sicily are by far more recent than Himalayan wolves, formerly considered a subspecies of *C. lupus* and named *C. l. lanigeror C. l. chanco*, but now ranked as distinct species *C. himalayensis* (Aggarwal et al. 2007). They diverged c. 550,000 (95% HDP = 495,100–605,600) years ago (Matsumura et al. 2014), and predated the evolutionary radiation of Eurasian and New World *C. lupus*.

In this study we used Matsumura et al. (2014) estimates of Japanese wolf mtDNA divergence time to compute a MRCA = 13,400 years (95% HPD = 4000 – 21,230) of the Italian wolf clade. The mtDNA genome of the Sicilian wolf is basal to the Italian wolf clade, thus a MRCA = 13,400 years can be considered as an approximate estimate of island-mainland DNA divergence. Although phylogenetic relationships and divergence time estimates obtained from complete mtDNA genomes should, in principle, outperform estimates obtained using only shorter sequences, they should anyway be used with caution. First, the mtDNA is a maternal haploid genome informative only on single-gene relationships and not on population-species phylogeny.Then, the sample size used in our and other studies (e.g., Angelici et al. 2018) are by far too small to exclude uncertainty. The wolf population in Sicily is extinct, the available museum specimens are few and perhaps not always suitable for genomic studies, thus the sample size of the target population could not be much expanded. However, we believe that the addition of complete mtDNA genomes from other haplogroup-2 wolf populations could improve the phylogenetic structure and connections of the Italian wolf clade, allowing more reliable estimates of divergence times. Moreover, sequences form chromosomal genes could contribute to better describe the phylogeographic history of the wolf population in Sicily.

The mtDNA CRs of Sicilian wolves are distinct from homologous CR sequences of historical Italian wolves obtained by Dufresnes et al. (2018). Wolves in peninsular Italy were certainly abundant a few centuries ago, but the museum specimens suitable to DNA sequencing are too few to conclude that the Sicilian haplotypes were absent in the historical mainland population. Hence, we cannot exclude that the Sicilian mtDNA haplotypes evolved in peninsular Italy. However, based on the available data, the most parsimonious hypothesis is that those haplotypes evolved in Sicily following wolf island colonization. The divergence time of 13,400 years (95% HPD = 4000 – 21,230) of the Sicilian wolf mtDNA is compatible with the age of the last land bridge between the island and the south-western tip of Italy, that is 21,500 – 20,000 (Antonioli et al 2012). A late Pleistocene colonization of peninsular wolves before the Messina strait was definitely flooded does not exclude earlier colonization waves, which, seems, nevertheless undocumented by the available Sicilian wolf specimens.

During late Holocene a number of species, and in particular ungulates (red deer, fallow deer, roe deer, wildboar), the natural prey of wolves, went extinct in Sicily like due to anthropogenic pressures (La Mantia and Cannella, 2008). The concomitant consequences of habitat transformations, ungulate decline and overhunting most probably pushed the wolf population of Sicily to decline and finally disappearing. The few available stuffed specimens evidence smaller body size and paler coat colours of the last wolves in Sicily in comparison to the Italian wolves. Moreover wolves in Sicily did not show the darker fur strip on the forearms, a peculiar morphological trait of the peninsular Italian wolf population (Altobello 1921; Ciucci and Boitani 2003). Dwarfism and local phenotypic adaptations are typical of some island vertebrate populations. Moreover, we cannot exclude that during the final population bottleneck wolves in Sicily crossbred and hybridized with free-ranging dogs, perhaps accelerating the speed of the extinction vortex (see: Gómez-Sánchez et al. 2018). Future genomic data set and analyses could perhaps shed more light on the extent of homozygosity and eventual domestic dog introgression in the lost populations of wolves in Sicily.

## Acknowledgements

We gratefully thank the Civic Museum “Baldassare Romano” (Termini Imerese) and dr. Fabio Lo Bono, curator of Mammals.

